# Persistent Underrepresentation of Women’s Science in High Profile Journals

**DOI:** 10.1101/275362

**Authors:** Yiqin Alicia Shen, Jason M. Webster, Yuichi Shoda, Ione Fine

## Abstract

Yiqin Alicia Shen, Jason M. Webster, Yuichi Shoda, and Ione Fine Department of Psychology, University of Washington Past research has demonstrated an under-representation of female editors and reviewers in top scientific journals, but less is known about the representation of women authors within original research articles. We collected research article publication records from 15 high-profile multidisciplinary and neuroscience journals for 2005-2017 and analyzed the representation of women over time, as well as its relationship with journal impact factor. We find that women authors have been persistently underrepresented in high-profile journals. This under-representation has persisted over more than a decade, with glacial improvement over time. Even within our limited group of high profile journals, the percent of female first and last authors is negatively associated with journal impact factor. Since publishing in high-profile journals is a gateway to academic success, this underrepresentation of women may contribute to the lack of women at the top of the academic ladder.

## Introduction

It has long been known that female representation within STEM fields decreases at every stage of the academic career (National Research Council, 2006; Valian, 1998). Take neuroscience as an example, in the year 2016, over 55% of graduate students were female, however, only 45% of postdoctoral researchers, and 32% of faculty were female (McKinley Advisors, 2017).

The reason behind such gender disparity is complex (McKinley Advisors, 2017; National Research Council, 2006; Valian, 1998). One potential problem that has gathered increasing research interest is gender discrepancies within high-profile scientific publications. For example, a series of prominent articles and editorials over the last decade have pointed out that women are underrepresented as authors of commissioned articles in *Nature* (Conley, 2005; "Gender imbalance in science journals is still pervasive," 2017; "Nature’s sexism," 2012; Shen, 2013).

While commissioned opinion pieces are important and changes in author recruiting can be directly influenced by journal policies, one of the most influential functions of high profile journals such as *Nature* is to disseminate original findings. An initial small-scale analysis to examine gender disparities in research articles across two 3-month periods in 2006 and 2016 in *Nature Neuroscience* found only a 1% increase in the number of female corresponding authors over that time period ("Promoting diversity in neuroscience," 2018).

In the current research, we extend this work by using data mining techniques to examine the proportion of female first and last authors for all research articles published between 2005 and 2017 across a wide range of high-profile journals that publish neuroscience research. In the results describe here, we focus on three findings: First we show that the proportion of women last authors in high profile research journals is much lower than the proportion of women scientists receiving USA RO1 grants or the European equivalents. Second, we show that, even within this highly selective group of journals, there is negative relationship between journal impact-factor and proportion of female first and last authors. Finally, we show that the lack of representation of female authors has remained dispiritingly unchanged in most journals over the last 13 years.

## Methods

The full details and code for data acquisition, processing, and analysis are provided in the Github Repo (https://github.com/VisCog/Women_in_high_profile_journals). Here we describe an overview of our approach.

### Data Acquisition

We downloaded metadata associated with all papers published from 2005 to 2017 from the PubMed’s MEDLINE database ("MEDLINE/PubMed Data," 2017). We then subset to focus on research articles in those journals by excluding articles without an abstract.

To focus on high profile journals, we selected 15 journals to include based on the 2016 impact factors from the Thomson Reuters InCite Journal Citation Report (Clarivate Analytics, 2016). Journals which focused on a particular aspect of neuroscience (e.g. EMBO, Stroke) were excluded. This resulted in a list that included both non-specialized multidisciplinary journals (Nature, Science, Proceedings of National Academy of Science (PNAS)), and top non-specialized journals in the field of neuroscience (Nature Review Neuroscience, Nature Neuroscience, Annual Review of Neuroscience, Behavioral and Brain Sciences, Neuron, Trends in Neurosciences, Brain, Cerebral Cortex, Neuropsychology Review, Current Opinion in Neurobiology, Journal of Neuroscience, NeuroImage). We then acquired the subset of the MEDLINE publication metadata based on this list of selected journals.

These steps resulted in a total of 166,979 records for those 15 top journals between the year 2005-2017 which were included for further analysis.

For comparison with our publication data, we also acquired data on the percentage of NIH RO1 grants in the U.S. and the percentage of MRC research grants in the U.K. awarded to women within this time period. This data was obtained from the NIH data book ("MEDLINE/PubMed Data," 2017) and MRC success rate data ("Medical Research Council 2016/17 Grant and Fellowship application success rates," 2018), respectively, in aggregated forms.

### Gender Determination

Due to the large quantities of publication records, manually classifying author gender is infeasible. Instead, we estimated author’s gender using genderizeR, a genderize.io interface for R (Wais, 2016). The genderize.io database currently contains 216286 distinct first names and gender self-report data from social media platforms across 79 countries and 89 languages. Based on each unique first name, it provides a gender prediction as well as a probability estimation for the prediction. We first conducted analysis with full set of data, then replicated our analysis with the subset of names for which gender assignment certainty was greater than 0.9.

### Analysis

To estimate the overall representation of women in each journal, we first calculated the overall percentage of female first and last author for each journal across the entire time range. To estimate the association between author gender ratios and journal profile, we calculated the Spearman’s rank order correlation between percentage of female first and last author with each journal’s 5-year impact factor (Clarivate Analytics, 2016), with or without the three multidisciplinary journals (i.e. Nature, Science, PNAS).

To see the trends of female representation over time, we also regressed the percentage of female first and last authors in each journal on time (measured in years). The resulting slopes are an indicator of the rate of change of female authorship in each journal.

## Results

The percentage of female authors for high-profile neuroscience journals is lower than expected based on the proportion of women scientists in the field. As shown in Figure 1, between 2005-2017, the percentage of female last authors are highest in Neuropsychology Review (39.04%) and Current Opinion in Neurobiology (27.19%), and were lowest in Nature (14.64%) and Science (15.53%). This pattern of results is similar for first authors, with Neuropsychology Review (52.58%) and Brain (43.01%) having the highest percentage of females, and Nature (25.22%) having the lowest. Also note that the percentage of female last authors in almost all journals (except for Neuropsychology Review) is lower than the percentage of females awarded prestigious grants such as NIH RO1 (∼30%, see grey line in Fig. 1 right panel), which is comparable to the proportion of UK Medical Research Council research awards. These grants are an important point of comparison since they are awarded based on the peer-evaluated qualities of significance, impact, research quality, and laboratory productivity.

**Figure 1.**
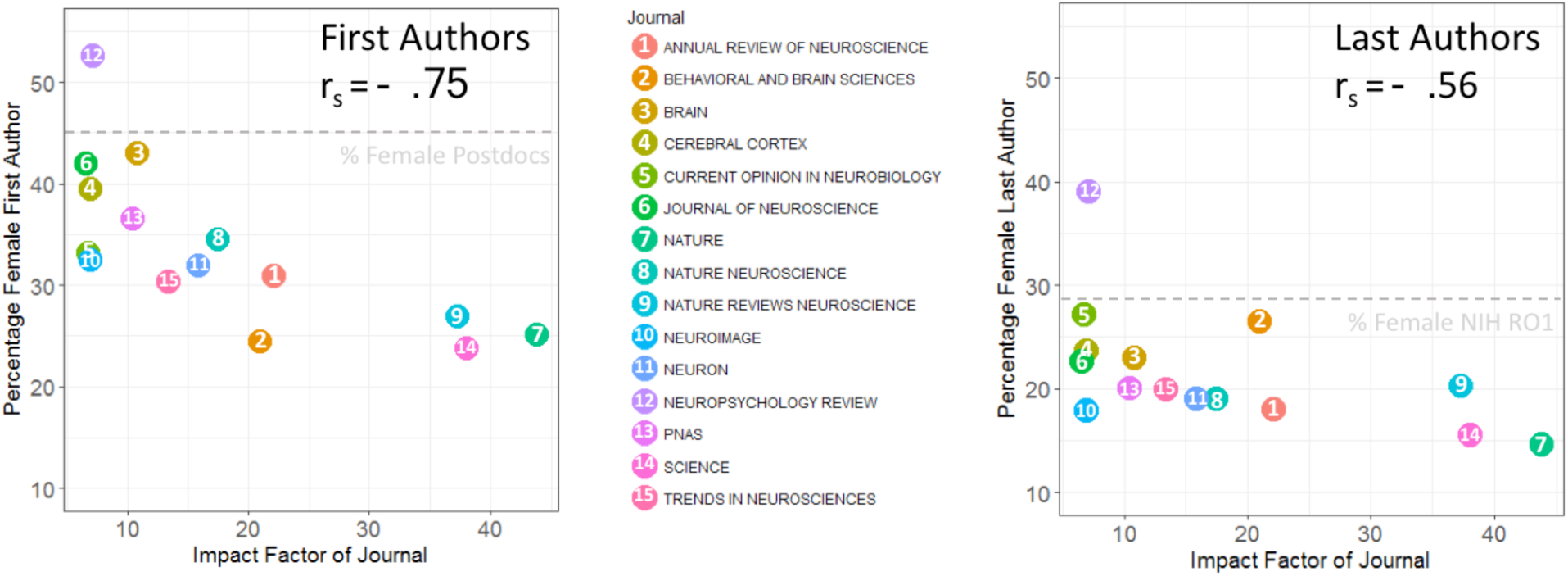
Percentage of female first and last authors between 2005-2017 vs. Journal’s 5-year impact factor.

Journals with higher impact factors have lower representation of female first and last authors. The percentage of both female first and last authors displayed a strongly negative association with journal impact factor (first author r_s_ = −0.75, *p* < .01, last author r_s_ = −0.56, *p* < .05). Even within the field of neuroscience, a higher impact is associated with a lower female representation: the same trend was found when excluding the multidisciplinary journals such as Nature, Science, and PNAS (first author r_s_ = −0.65, *p* < .05, last author r_s_ = −0.32, *ns*).

As shown in Figure 2 and Table 1, the percentage of women first and last authors has increased at less than 1% per year for almost all journals. However, there are variations between journals. Some journals, such as Brain, have a steady increase of female last authors of over 1% per year, while other journals, such as Nature Neuroscience, have a decrease in the percentage of female last authors per year. Over that period, the percentage of women receiving NIH RO1 awards has remained roughly constant at ∼30%, a percentage that reflects the number of women at the Associate/Professor level within STEM fields (see grey line in Figure 2B).

**Figure 2.**
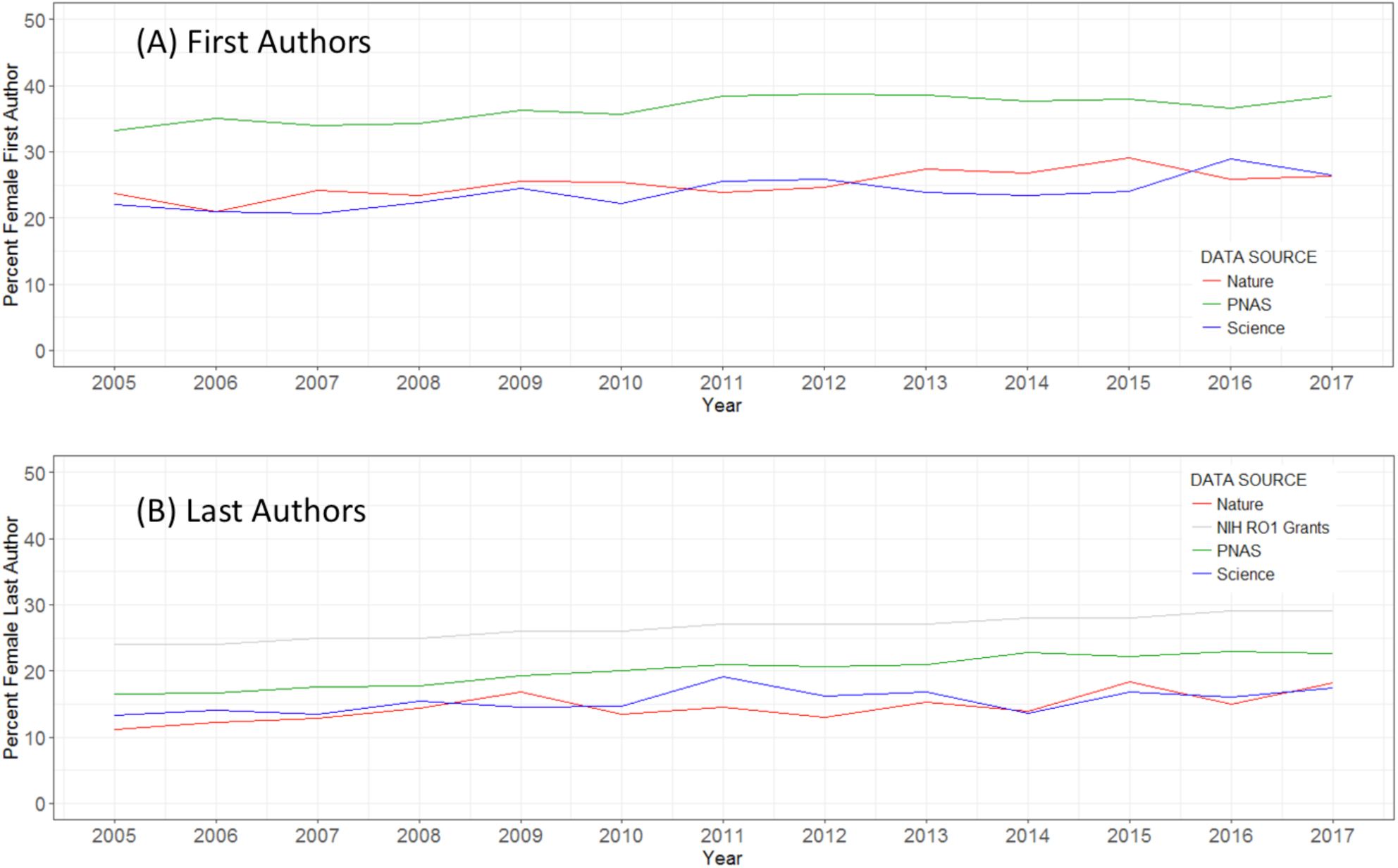
Change in percentage of female first (Panel A) and last (Panel B) authors overtime.

**Table 1.** Percentage change for female first and last authors per year.

| **Journal** | **% Change in Female First Author Per Year (Slope)** | **% Change in Female Last Author Per Year (Slope)** |
| --- | --- | --- |
| Neuropsychology Review | 1.85 | -0.36 |
| Nature Neuroscience | 0.20 | -0.11 |
| Nature Reviews Neuroscience | 0.97 | 0.08 |
| Trends in Neurosciences | 0.59 | 0.19 |
| Neuron | 0.23 | 0.22 |
| Annual Review of Neuroscience | 0.98 | 0.22 |
| Science | 0.47 | 0.27 |
| Current Opinion in Neurobiology | 0.22 | 0.31 |
| Nature | 0.40 | 0.40 |
| Behavioral and Brain Sciences | 1.80 | 0.44 |
| Journal of Neuroscience | 0.60 | 0.53 |
| PNAS | 0.40 | 0.58 |
| NeuroImage | 0.86 | 0.77 |
| Cerebral Cortex | 0.76 | 0.78 |
| Brain | 1.42 | 1.03 |

Analysis restricted to author names with gender assignment certainty of greater than 90% produced qualitatively identical results (data not shown).

## Discussion

Using data mining techniques, we evaluated the publication records of original research articles for the top 15 journals publishing neuroscience research from 2005-2017. We found that 1) proportion of women authors in high profile research journals is substantially lower than the proportion of women receiving competitive grants, 2) there was a negative relationship between journal impact-factor and proportion of female first and last authors, and 3) the rate of increase in female representation is on average less than one percent per year for first authors and less than half a percent per year for last authors.

While our research clearly demonstrated gender discrepancies, our data does not speak to the underlying causes. One possibility is that women are submitting less to high profile journals. Editors of *Nature Neuroscience* reported only 21.5% of submissions to their journal were from female authors ("Promoting diversity in neuroscience," 2018). Another possibility is that women are less successful in negotiating prestigious authorship positions. While women are more likely to be the person performing experiments (Macaluso, Lariviere, Sugimoto, & Sugimoto, 2016), they are less likely to be in the prestigious lead author positions (West, Jacquet, King, Correll, & Bergstrom, 2013). A third possibility is bias in the publication pipeline. Experimental evidence suggests that, when reviewers are randomly assigned to evaluate scientific work ostensibly submitted by a female or a male author, they rated the work written by male authors as having higher rigor (Knobloch-Westerwick, Glynn, & Huge, 2013). Nature, in a series of editorials spanning more than a decade, also observed that its editors are less likely to ask women to write commissioned pieces ("Nature’s sexism," 2012). Clearly, more research is needed to evaluate the relative importance of those underlying mechanisms.

Like it or not, publication in high-profile journals remains an important gateway for career advancement. High-profile publications have an enormous impact on the likelihood of receiving awards, funding, and positions in highly ranked research institutions. Conversely, the lack of high-profile publications may partially account for the lower rate of recruitment, retention, and promotion for women faculty (McKinley Advisors, 2017; National Research Council, 2006; Shen, 2013; Valian, 1998). The under-representation of women in high profile journals impacts thousands of talented scientists.

It is now well past time for high-impact journals to begin collaborating with the scientific community to develop and validate evidence-based procedures to remove sources of bias throughout both the editorial and the reviewing process for original scientific articles. We would recommend some obvious first steps. First, all journals should collect gender and minority statistics on submission and acceptance rates for papers and should make these data publically available. Second, journals should use mandatory double-blind reviewing. Results from other disciplines suggest that double-blind reviewing procedures significantly increase the proportion of female lead research articles (Budden et al., 2008). Finally, reviewers should be provided with clearer guidance about review criteria, as is done for NIH review panels.

## Acknowledgements

We thank Dr. Tara M. Madhyastha, Dr. Ariel Rokem, and Kelly Chang for their help with code review and maintenance.

